# Using camera traps to determine occupancy and breeding in burrowing seabirds

**DOI:** 10.1101/2021.04.08.438875

**Authors:** Jeremy P. Bird, Richard A. Fuller, Penny P. Pascoe, Justine D. S. Shaw

## Abstract

Burrowing seabirds are important in commercial, ecological and conservation terms. Many populations are in flux owing to both negative and positive anthropogenic impacts, but their ecology makes measuring changes difficult. Reliably recording key metrics, the proportion of burrows with breeding pairs, and the success of breeding attempts, requires burrow-level information on occupancy. We investigated the use of camera traps positioned at burrow entrances for determining the number of breeding pairs in a sample to inform population estimates, and for recording breeding success. Linear Discriminant Analyses of time series activity patterns from camera traps successfully partitioned breeding and non-breeding burrows at different stages of the breeding season and had reasonable predictive ability to determine breeding status on a small test dataset. Compared with traditional techniques for determining burrow occupancy (e.g. manual burrow inspection and playback of conspecific calls at burrow entrances), camera traps can reduce uncertainty in estimated breeding success and potentially breeding status of burrows. Significant up-front investment is required in terms of equipment and human resources but for long-term studies camera traps can deliver advantages, particularly when unanticipated novel observations and the potential for calibrating traditional methods with cameras are factored in.

## INTRODUCTION

Seabirds are a major component of marine food webs consuming prey volumes commensurate with commercial fisheries (Barrett et al., 2006; Danckwerts et al., 2014). In the Southern Oceans one taxonomic group, the Procellariiform petrels, is second only to penguins in terms of avian biomass (Catry et al., 2003) and volume of prey consumed (Croxall et al., 1984). By transferring nutrients from pelagic to marine and coastal ecosystems, petrels make significant contributions to global geochemical cycles (Otero et al., 2018). These inputs, together with soil modification from their burrow-nesting habit result in positive influences on island biodiversity (Graham et al., 2018; Sekercioglu, 2006). However, petrels are also highly susceptible to invasive predators which have precipitated catastrophic declines in many populations resulting in 45 of 97 species (46%) being threatened with extinction (BirdLife International, 2018; Croxall et al., 2012; Dias et al., 2019).

Collectively there are strong commercial, ecological and conservation incentives for understanding petrel population trends, but they are difficult to monitor (Rodríguez et al., 2019). Seasonal occurrence, nocturnal activity at colonies and challenges in finding, counting and ascertaining the contents of burrows all contribute to high levels of uncertainty when populations are assessed (Bird et al., submitted; Rodríguez et al., 2019). The most important metrics for tracking conservation status and population-level responses to impacts are the number of breeding burrows and their success rate, both of which rely upon one-off, or repeat measures of burrow occupancy.

In recent years, audio playback for small species and manual burrow inspections for larger species have emerged as the most widely used occupancy detection methods in burrowing seabird studies (Bird et al., submitted). Playback requires calibration to account for non-responsive birds (Soanes et al., 2012). This is an invasive and time-consuming process. Because response rates vary seasonally and between life history stages (Berrow, 2000; Ratcliffe et al., 1998; Ryan et al., 2006) repeated calibration would be required to collect population-level information on breeding success. This is unethical and logistically impractical. Furthermore, detailed ecological study at the burrow level using playback is not possible because response rates of individual birds vary temporally. As such, playback is only suitable for determining the proportion of occupied burrows at a population-level (Parker and Rexer-Huber, 2016). Studies typically assume that occupied burrows represent breeding pairs, and the occupancy rate is then used to adjust burrow counts when estimating breeding population size. Yet not all occupied burrows are occupied by breeding birds, resulting in unquantified error in final estimates (Parker and Rexer-Huber, 2016). In contrast, burrow inspection can provide more detailed information about individual burrow contents, but it is disruptive to petrels. Investigator disturbance can result in high rates of nest abandonment, reduced hatching success, high divorce rates between disturbed pairs, and emigration of immature birds from natal colonies (Carey, 2009). To meet ethical guidelines and ensure data is representative of natural parameters, manual burrow inspection has evolved away from early methods which involved partial burrow excavation. In recent times, the use of purpose-built burrow inspection scopes has become prevalent, but accuracy is highly variable according to the quality of equipment available (Carlile et al., 2019; Lavers et al., 2019). Manual burrow inspection is also labour intensive, and repeat visits to measure breeding success still have uncertainty around outcomes when burrows change from occupied to empty between visits.

Use of camera traps in ecological studies has become common because they can provide time series data on the activity of individuals (Wearn and Glover-Kapfer, 2019). They have been used to measure seabird breeding phenology and provide indices of abundance for above-ground nesting species, but their use in studies of burrowing seabirds has mainly been limited to understanding predation pressures by invasive alien species (Edney and Wood, 2020). Fischer et al. (2017) assessed the suitability of camera traps for studying diving-petrels *Pelecanoides* spp. by pairing them with Radio Frequency Identification tags deployed on five birds, with tag readers at the burrow entrance. They found detection rates were low, perhaps owing to the size and behaviour of the birds—diving-petrels are small, arrive and depart rapidly from their burrows, and are very well insulated reducing the chance of infra-red triggers—and concluded camera traps appear unsuitable for studying breeding biology in diving-petrels.

There has been no test of whether continuous observations using camera traps can overcome shortcomings of playback and manual burrow inspection. Unlike playback no calibration is required, and long-term deployments could reveal outcomes of breeding events in individual burrows with higher resolution than intermittent manual inspections. If so, camera traps have the potential to considerably improve burrowing seabird conservation assessments providing reliable information with low uncertainty on occupancy, the proportion of burrows occupied by breeding pairs, and breeding success.

Here we investigate the use of camera traps for monitoring the breeding biology of several species of burrowing petrel on sub-Antarctic Macquarie Island. We assess the performance of cameras for: i) recording nest-level activity; ii) recording breeding success rate; and iii) inferring the ratio of breeding to non-breeding burrows. Finally, we compare the results from camera traps with traditional playback and burrow-inspection methods.

## METHODS

### Study site and species

Macquarie Island (54°30‘S, 158°57’E) has many of the characteristics that make surveying and monitoring burrowing petrels challenging. Lying approximately 1,500 km south-east of Tasmania, almost midway between Australia and Antarctica, it is remote and therefore costly to visit. The island’s status as a UNESCO World Heritage Site and Tasmanian Nature Reserve means invasive research activities and visitation are restricted. It is 12,785 ha in area, 34 km long and up to 5 km wide, rugged with sections of coastal cliffs and a steep escarpment rising to an upland plateau reaching 400 m above sea level. The sub-Antarctic climate and potential for extreme weather conditions (Adams, 2009; Pendlebury and Barnes-Keoghan, 2007), the terrain, and the limited access to medical facilities mean strict health and safety protocols are in place to protect personnel, and these also place constraints on fieldwork capacity. Furthermore, the island’s size, terrain, steep slopes and fact that vegetation communities are recovering following decades of impacts by European Rabbits *Oryctolagus cuniculus* (Springer, 2016) all make finding and studying seabird burrows difficult. The friable peat soils in many seabird colonies are fragile. Safe, low impact access is challenging.

Our study addressed two large-bodied species: Grey Petrel *Procellaria cinerea* (950-1,220 g) and White-headed Petrel *Pterodroma lessonii* (750-780 g); and two medium-bodied species: Blue Petrel *Halobaena caerulea* (171-253 g) and Antarctic Prion *Pachyptila desolata* (116-160 g; Jouventin, 1985). All four species nest in burrows dug predominately in peat substrate.

Constraints on the location and timing of fieldwork meant we were unable to compare all burrow assessment methods across all species, but we compare the utility of each method at least between a large-bodied and a medium-bodied species.

### Camera-traps

Six Reconyx HC600 trail cameras were deployed at the entrance to Grey Petrel burrows from 10^th^ June 2017 to 22^nd^ November 2017. In 2018, six Reconyx HC600 and four Spypoint Force 10 trail cameras were deployed outside Grey Petrel burrows from 1^st^ April to 2^nd^ November (Figure 1). Twenty Spypoint Force 10 cameras were deployed outside Blue Petrel burrows from 15^th^ August 2018 to 4^th^ February 2019 (see Figure S1-5 for burrow identification). Cameras were all set to ‘high’ sensitivity, to take three images per trigger with 1 second interval between, and no delay between triggers. They were checked at intervals to replace batteries and memory cards.

**Figure 1:**
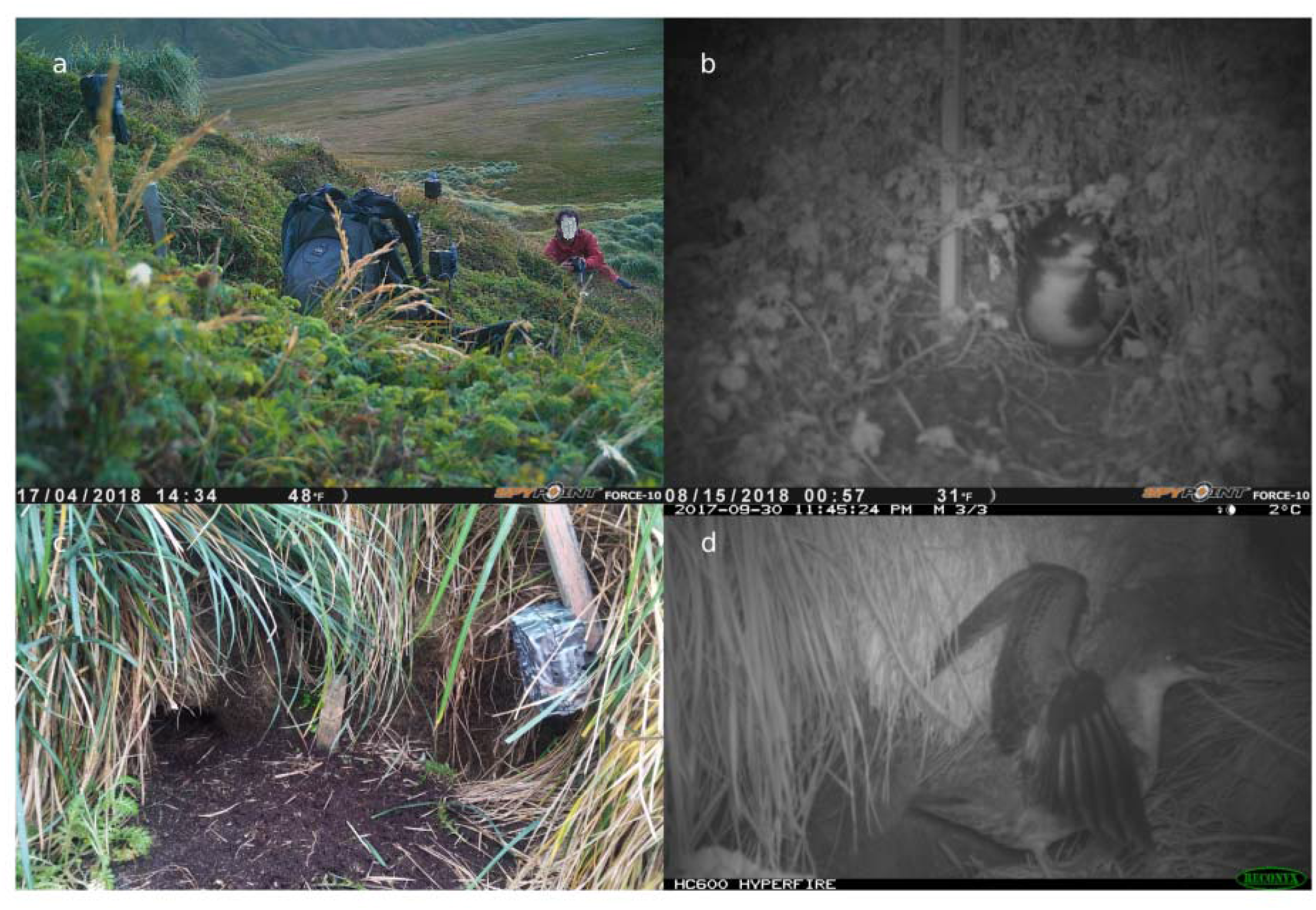
Camera trap set-up and results: a) checking Spypoint cameras in a Blue Petrel colony, b) a Blue Petrel emerging from its burrow, c) a Recconyx camera outside a Grey Petrel burrow, and d) a Grey Petrel chick close to fledging exercising outside its burrow.

We tagged all images manually in ExifPro 2.1.0 (http://www.exifpro.com/index.html), discarding any images associated with camera set-up. We tagged images that featured a bird with the species, the number of individuals (rarely >1), and whether there was a chick. We also tagged behaviours: burrow entries, exits and tidying of the burrow entrance which appeared to be associated with breeding.

To assess camera performance, we recorded periods of camera inactivity, we compared the total numbers and ratios of false triggers to target triggers, the mean numbers of events per day between cameras, and the ratio of entries to exits. For behavioural analysis, we grouped images into ‘events’—series of consecutive images separated by no more than 5 minutes. We plotted the number of events per day against date for both adults and chicks. We relabelled entries and exits as ‘transits’ and plotted the total per day against date, together with the number of tidying events recorded.

We manually inspected all Grey Petrel burrows with camera traps at the entrance four times during the period over which cameras were deployed (5 months in 2017 and 7 months in 2018) to determine breeding status and breeding success at the burrow. To understand whether adult activity patterns derived from the cameras could be used to reliably infer breeding status of burrows we derived test variables: mean number of events, duration of absences in days, the proportion of days where high activity was recorded (> 5 events), and the proportion of days where low activity was recorded (1-5 events). We derived these variables for each stage of the breeding season: incubation, early chick-rearing and late chick-rearing (fledging was dropped owing to low adult activity). We tested for correlation between the variables by dropping one of a pair of variables if their *r*^2^ was > 0.6, and then used principal component analysis (PCA) to look for patterns in our data. Then we classified breeding and non-breeding burrows using linear discriminant analysis (LDA). PCAs and LDAs were performed for each breeding stage. Because our burrow sample was small, we used all full-season time series to build the LDA models. We derived a test dataset from cameras that malfunctioned during the season for which only partial time-series were available and used these to test the models. Breeding success was derived from chick activity recorded at the burrow entrance. Analyses were conducted in R using the ‘tidyverse’, ‘MASS’, ‘FactoMineR’ and ‘caret’ packages (Kuhn, 2008; Lê et al., 2008; R Core Team, 2020; Venables and Ripley, 2002; Wickham et al., 2019, see Supplementary Code).

### Playback and burrow inspection

Both playback and manual burrow inspection have been used to determine occupancy of Grey Petrel, Antarctic Prion and Blue Petrel burrows (Barbraud et al., 2009; Brothers and Bone, 2008; Dilley et al., 2017). However, *Pterodroma* petrels vocalise during aerial displays at night, but rarely respond from the nest chamber to stimuli during the day, so manual inspection is required for White-headed Petrels (Brothers, 1984; Sinclair, 1981). We preferentially chose to manually inspect burrows because we were concerned that variable response rates to playback would affect nest-level data (Schulz et al., 2006), but we compared with playback for medium-bodied species where we had low confidence in manual inspections.

We first visited burrows during incubation or early chick-rearing in the 2017-2018 breeding seasons of our four study species. We inspected 237 White-headed Petrel burrows with a homemade burrowscope (Figure S6). A second observer independently checked the same burrows with a handheld Canon Powershot camera held at arm’s length down the burrow to calibrate the two methods and allow correction of data from previous years when cameras were used. We checked 242 Grey Petrel burrows by handheld camera or burrowscope, and 247 Blue Petrel and 76 Antarctic Prion burrows with the burrowscope. A burrow was recorded as ‘empty’ when the observer believed they could see the end of the burrow and any side chambers were visible in images, or ‘uncertain’ if the end was not visible. In occupied burrows we noted the number and identity of adult birds, eggs, and chicks.

We revisited burrows to check breeding status and the outcome of breeding attempts through the season. All burrows in which eggs, recent eggshell or chicks were found in the burrow or at the entrance were recorded as breeding burrows. If adults were present during the first visit but not subsequently the breeding status was ‘uncertain’. Burrows with adults present later in the season were recorded as non-breeding. The repeat visits were used to calibrate earlier visits—the status of burrows labelled originally as ‘uncertain’ often became clear on second or third visits, and we estimated our type 1 error rate by identifying breeding burrows originally labelled as ‘empty’.

For comparison with occupancy and uncertainty estimates from our initial burrow inspections we used playback for both medium-sized species. Playback provides information on the proportion of burrows occupied, relying on birds responding when a recording of a conspecific vocalising is played near their burrow entrance. It is common in studies of small-bodied species whose burrow entrances are small, precluding manual inspection (Perkins et al., 2018). Response rates are known to vary so following Soanes et al. (2012) we repeated playback 4 times to a sample of 61 Antarctic Prion burrows and 9 times to 66 Blue Petrel burrows and used the ‘du Feu’ single-session mark/recapture method of population estimation to estimate the overall proportion of occupied burrows in our sample with confidence limits (du Feu et al., 1983).

## Results

### Recording nest-level activity and breeding success from cameras

Overall camera traps proved to be very effective. The expected target species was recorded on all cameras placed at burrow entrances. One camera placed at a burrow entrance for a night equates to one ‘trap night’. We recorded 47,179 images of Blue Petrels during 3,554 trap nights, and 42,705 images of Grey Petrels during 2,677 trap nights. However, performance varied greatly between the two types of cameras we deployed. The ratio of target triggers to false triggers was much higher for Reconyx cameras compared with the SpyPoint cameras, and the proportion of trap nights where >0 images were captured, i.e. the camera was known to be recording, was also higher (Table 1). In reality, short periods of several days where no images were captured are likely to reflect genuine absences of activity, whereas long periods (e.g. >10 days) with no target or false triggers likely reflect camera malfunction, especially in SpyPoint cameras given their high false trigger rate. Although we primarily deployed Spypoint cameras at Blue Petrel burrows and Reconyx at Grey Petrel burrows, the relative differences in performance are still apparent in the one Reconyx camera at a Blue Petrel burrow and the four SpyPoint cameras at Grey Petrel burrows (Table 1). Overall SpyPoint cameras were far more likely to be triggered when no animal was present, and there were many more days, irrespective of species, when the cameras did not record any images.

**Table 1:** Performance metrics for Reconyx and SpyPoint cameras positioned at seabird burrow entrances.

| make | species | n | target:false | proportion of trap | entries:exits |  |  |
| --- | --- | --- | --- | --- | --- | --- | --- |
|  |  | burrows | triggers | nights >0 images | min | max | SD |
| Reconyx | All | 13 | 18.86 | 0.64 | 0.20 | 1.56 | 0.41 |
| Reconyx | Grey Petrel | 12 | 19.89 | 0.63 | 0.20 | 1.56 | 0.41 |
| Reconyx | Blue Petrel | 1 | 6.39 | 0.85 | 0.96 | 0.96 | 0.00 |
| SpyPoint | All | 24 | 1.30 | 0.39 | 0.52 | 5.67 | 1.09 |
| SpyPoint | Grey Petrel | 4 | 3.08 | 0.33 | 0.60 | 1.07 | 0.17 |
| SpyPoint | Blue Petrel | 20 | 0.95 | 0.40 | 0.52 | 5.67 | 1.16 |

Despite the stark difference in camera performance, we recorded specific behaviours at all burrows such as entries and exits, digging, tidying at burrow entrances, chicks emerging and exercising, predators (Brown Skua *Catharacta antarctica*) and competitors entering burrows (e.g. Antarctic Prions in Blue Petrel burrows and Sooty Shearwaters in Grey Petrel burrows; Figures S7-8). While we were interested in burrow-level variation in behaviours recorded, the variance in certain behaviours, for example the ratio of entries to exits, suggests that positioning of cameras has an important bearing on what is recorded. Some cameras recorded five times more entries than exits, while others recorded five times more exits than entries (Table 1).

While camera make and positioning influenced the type and level of activity recorded at each burrow, averaging activity by classifying events and specific behaviours, both makes of camera provided season-long activity signatures for many burrows (Figures 2 and 3). The mean number of events recorded per day that burrows were visited was 4.10 for Blue Petrels and 2.58 for Grey Petrels. For both species the between-burrow variance in the mean number of events per day (SD 1.73 for Blue Petrels and SD 1.12 for Grey Petrels) was lower than the within-burrow variance (SD 3.95 for Blue Petrels and SD 1.76 for Grey Petrels), suggesting that the activity captured was relatively uniform across burrows.

**Figure 2:**
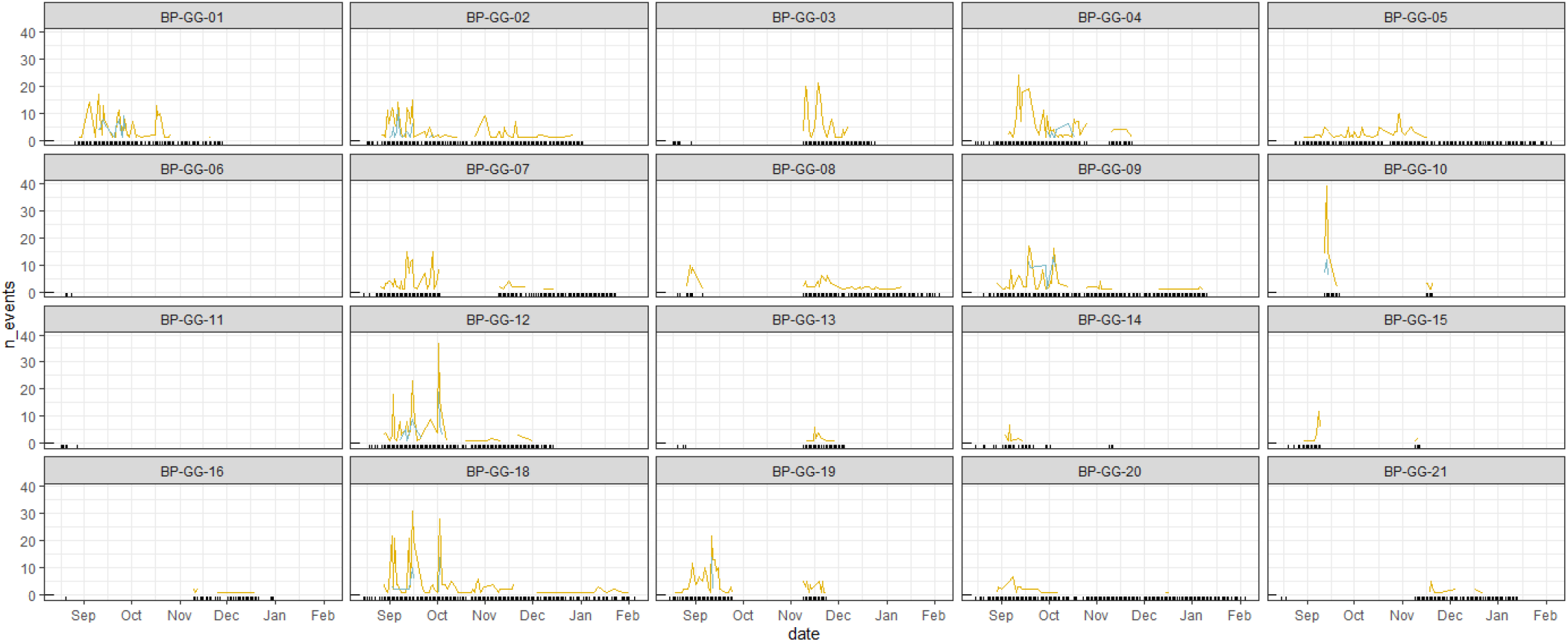
Combined entries and exits (yellow) and tidying events (blue) recorded at Blue Petrel burrow entrances during the breeding season. Rug plots show the days on which any images were captured, i.e. the camera was known to be functioning correctly.

**Figure 3:**
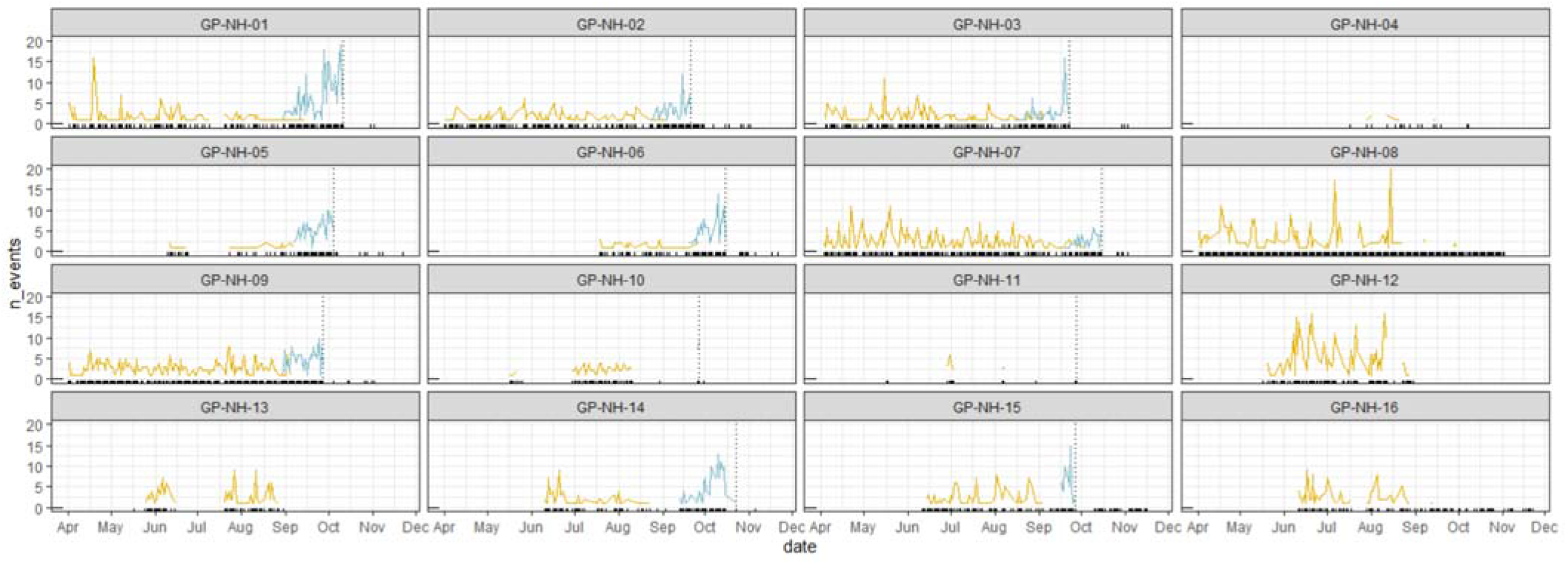
Total number of events featuring adults (yellow) and chicks (blue) at Grey Petrel burrow entrances during the breeding season. Rug plots show the days on which any images were captured, i.e. the camera was known to be functioning correctly. Vertical dotted lines represent inferred fledging dates.

Peak activity at Blue Petrel burrows is apparent early in the breeding season, tailing off later in the season (Figure 2). There are cyclic patterns in activity spikes with periods of absence or reduced activity between them. Generally, we also observed bouts of tidying around the burrow entrance early in the breeding season at burrows with high activity peaks. Burrows 05 and 20 lack early-season activity peaks, and burrow activity ceases earlier in the season than at e.g. burrows 02, 09 and 18. Unfortunately it is also evident from the rug plots in Figure 2 how poorly the Spypoint cameras performed (burrows 02-21) with few cameras recording right through to the end of the season.

The Grey Petrel dataset was more complete, with the exception of the Spypoint cameras at burrows 10, 11 and 13 (Figure 3). Again, there are clear cyclic patterns of burrow attendance. Recording began during incubation by which point adult attendance at most burrows was relatively uniform. During nest inspections no eggs, chicks or signs of breeding such as nest material were recorded in burrows 04, 08, 12, 13 or 16, and each of these burrows was only occupied during the first visit. These burrows were empty during all subsequent visits and were all considered to be non-breeding burrows after multiple visits. Nevertheless, the cameras show that all of them remained in use for much of the season. Burrows 08 and 12 have higher activity peaks than the breeding burrows at comparable stages of the breeding season, and the gaps between peaks appear to be longer.

Chicks were recorded in all other burrows during manual inspections, but the fate of the breeding attempts was sometimes uncertain. For example, burrow 03 was empty when visited on 26^th^ October but the camera showed the chick had first emerged when downy some weeks earlier, and had attained full adult plumage when last recorded on the camera on 23^rd^ October. From the cameras we inferred that all chicks in these study burrows fledged successfully, as well as the fledging date from each burrow (Figure 3).

### Inferring breeding and non-breeding burrows

We were particularly interested in whether the activity signatures from each burrow can be used to infer breeding status of the burrows. From the PCA there is weak evidence that absences tend to be longer during incubation and there are more low-activity days at breeding Grey Petrel burrows than non-breeding burrows (Figure 4). Later in the season during early and late chick-rearing absences are longer at non-breeding than breeding burrows, but when birds are present there is more activity at the burrow entrance of non-breeding burrows while breeding burrows are characterised by low activity (Figure 4).

**Figure 4:**
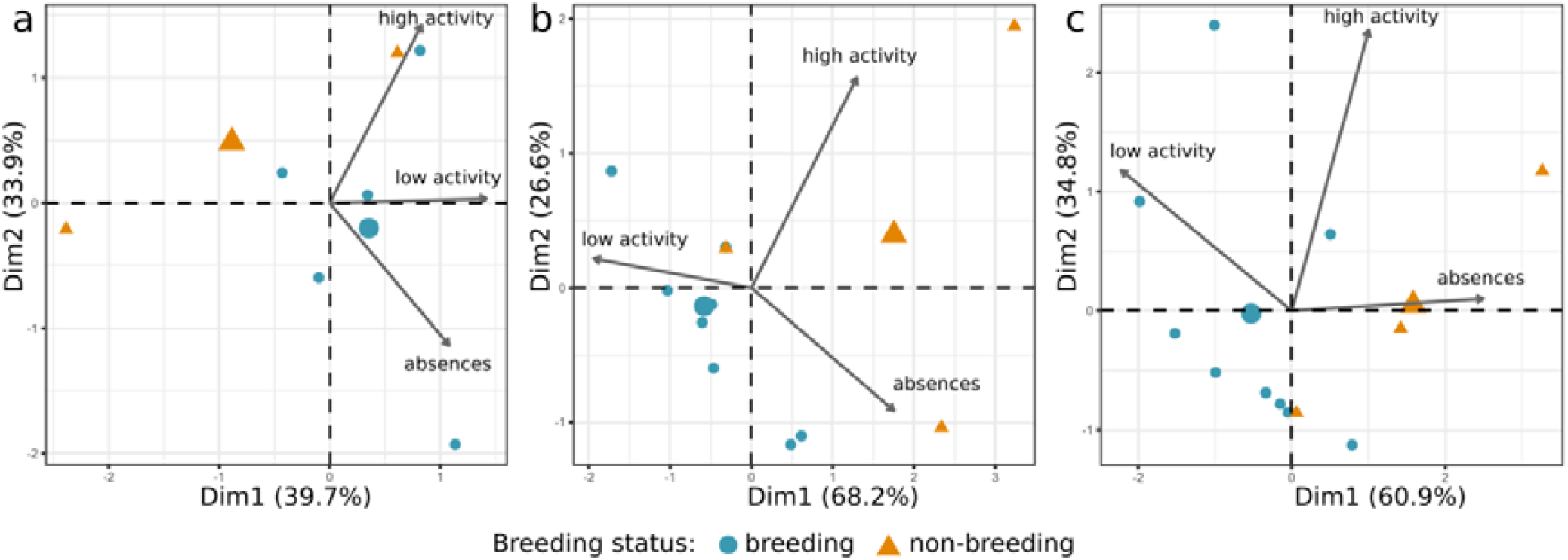
Principal Component Analyses of variables derived from adult activity data captured by cameras at Grey Petrel burrow entrances. Plot a) is activity data from during incubation, b) is from early chick-rearing and c) is from late chick-rearing. The larger symbols represent group means.

These apparent differences are also evident in the group means of the predictor variables we used in LDAs (Table 2). The three LDAs were broadly able to discriminate between breeding and non-breeding burrows in our training data (Figure 5). However, the analyses of absences and activity data during incubation and late chick-rearing misclassified the non-breeding burrow 13 based upon the partial time series available (Table 3).

**Table 2:** Group means for variables derived from adult activity at Grey Petrel burrow entrances that were used in Linear Discriminant Analysis, and their contributions to the first linear discriminant.

| stage | variable | breeding | non-breeding | LD1 |
| --- | --- | --- | --- | --- |
| incubation | absences | 1.87 | 1.70 | -0.09 |
|  | high activity | 0.05 | 0.10 | 13.63 |
|  | low activity | 0.59 | 0.28 | -12.30 |
| early chick-rearing | absences | 1.89 | 2.55 | 0.71 |
|  | high activity | 0.03 | 0.18 | 9.32 |
|  | low activity | 0.59 | 0.38 | -2.65 |
| late chick-rearing | absences | 3.32 | 6.75 | 1.05 |
|  | high activity | 0.02 | 0.04 | -11.69 |
|  | low activity | 0.34 | 0.20 | 5.53 |

**Table 3:** Predicted breeding status in two known-status test burrows from Linear Discriminant Analyses.

|  |  | prediction |  |
| --- | --- | --- | --- |
|  | LDA model | breeding | non-breeding |
| **GP-NH-10:**<br>**breeding** | incubation | - | - |
|  | early chick-rearing | 0.9995 | 0.0005 |
|  | late-chick-rearing | 0.9986 | 0.0014 |
| **GP-NH-13:**<br>**non-breeding** | incubation | 1.0000 | <0.0001 |
|  | early chick-rearing | 0.0911 | 0.9089 |
|  | late-chick-rearing | 0.9998 | 0.0002 |

**Figure 5:**
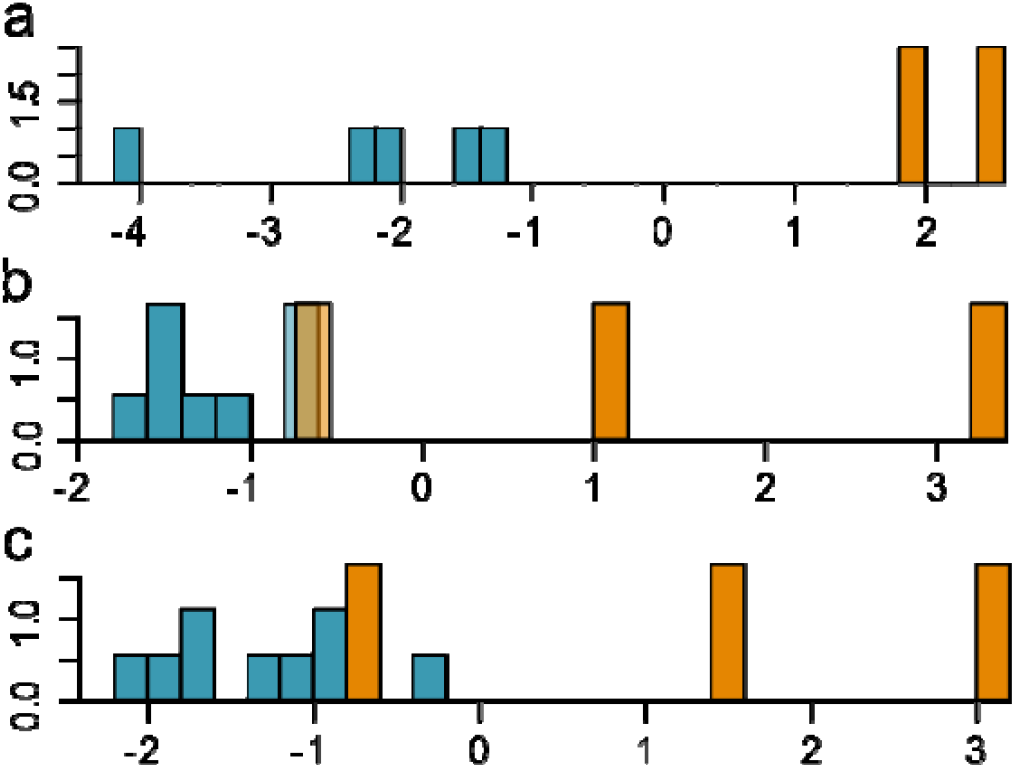
Linear Discriminant Analyses of breeding and non-breeding Grey Petrel burrows based upon variables derived from adult activity at burrow entrances. Model a) is activity data from during incubation, b) is from early chick-rearing and c) is from late chick-rearing. Breeding burrows are depicted in blue, non-breeding in orange.

### Comparisons with traditional methods

First, we compared approaches for measuring burrow occupancy. Figure 6 shows that using a burrowscope resulted in higher sensitivity (true positives) and specificity (true negatives), lower uncertainty and lower type 1 error (false negatives) when predicting burrow occupancy compared with using playback. For example, comparing panel a) using a burrowscope to inspect Antarctic Prion burrows with panel d) using playback, of 76 burrows inspected with a burrowscope we correctly assigned 20 burrows as empty and 15 as occupied on the first visit—sensitivity + specificity = 46%. However, we incorrectly identified 11 burrows as empty on first visit which were in fact occupied— type 1 error = 24%. And we were uncertain about the occupancy of 30 burrows (39%) on first visit, 14 of which were subsequently identified as occupied while 16 remained uncertain. Using playback, owing to our inability to distinguish between non-responsive birds in burrows and empty burrows, we could not definitively assign any burrows as empty (Figure 6d). We correctly identified 72 burrows as occupied on first visit—sensitivity 26%—but were uncertain about the status of the remaining 197 burrows (74%).

**Figure 6:**
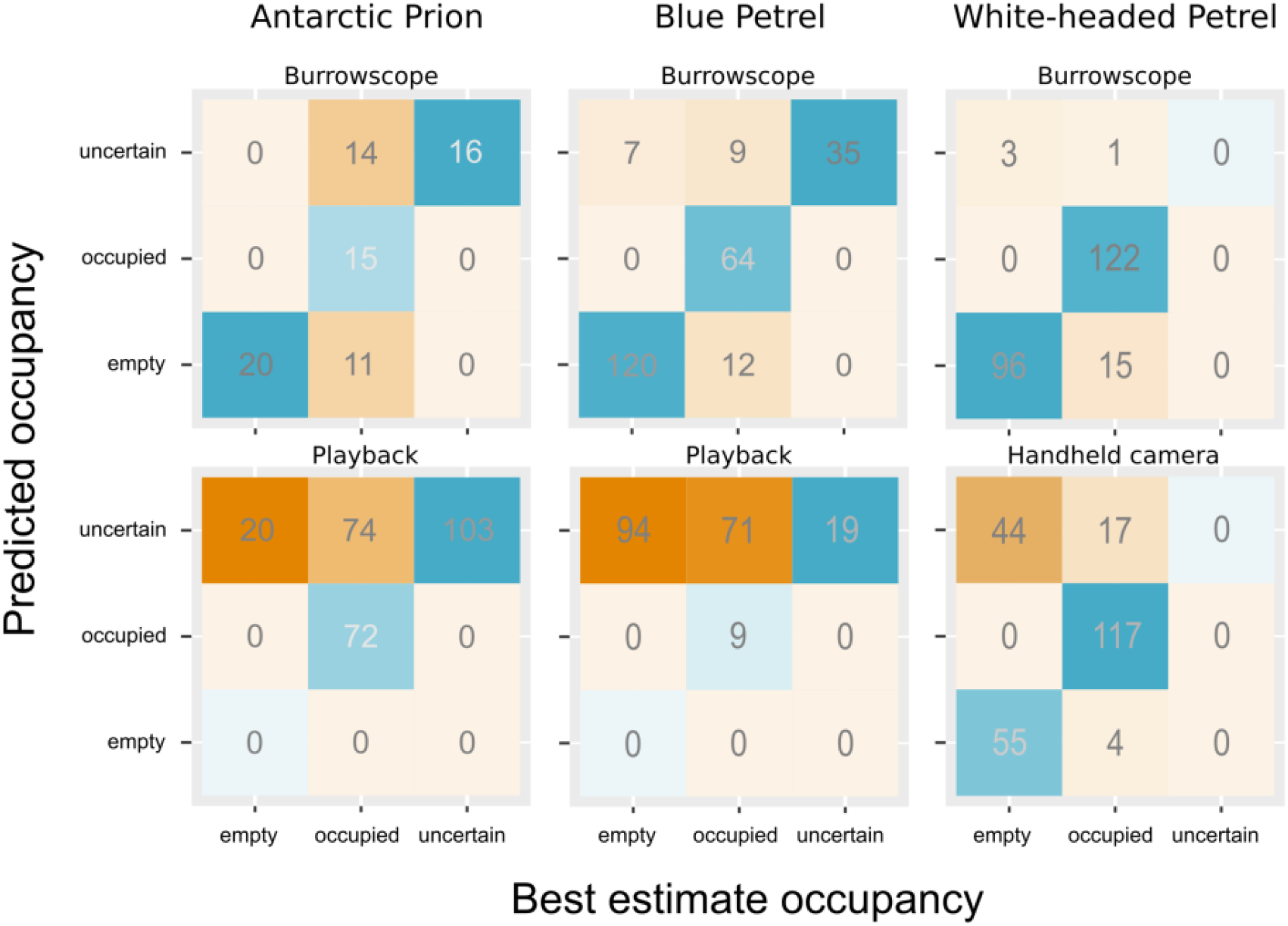
Confusion matrices of predicted versus observed occupancy from different methods. Predicted occupancy is based upon the first visit only, equivalent to one-off occupancy estimation feasible in most surveys. Observed occupancy is based upon all methods and all visits. Predicted = observed is shaded green, predicted ≠ observed is shaded yellow, with higher matching or mismatching ratios shaded darker.

There is an important distinction between determining occupancy in individual burrows and estimating overall population-level occupancy. The former is used to measure burrow-specific variables such as breeding status and success. The latter to adjust burrow counts when estimating the number of occupied burrows, either in a sample or in the whole population. We were able to refine population-level estimates of occupancy derived from playback by calibrating the response rate through repeat visits. We estimated overall occupancy in Blue Petrel burrows as 0.56 (95% CI: 0.42-0.70) after nine visits to calibrate our sample (Table S1). However, after achieving only 4 repeats for Antarctic Prions there was little flattening of the accumulation curve and the “du Feu” method, possibly erroneously in this case, estimates that all burrows are occupied (Table 4). Through repeated playback population-level occupancy can be estimated. However, while using a burrowscope is challenging for medium-bodied species owing to the small size of their burrow entrances, it still yielded tighter confidence intervals around the estimated occupancy statistic than repeat-measures playback (Table 4).

**Table 4:**
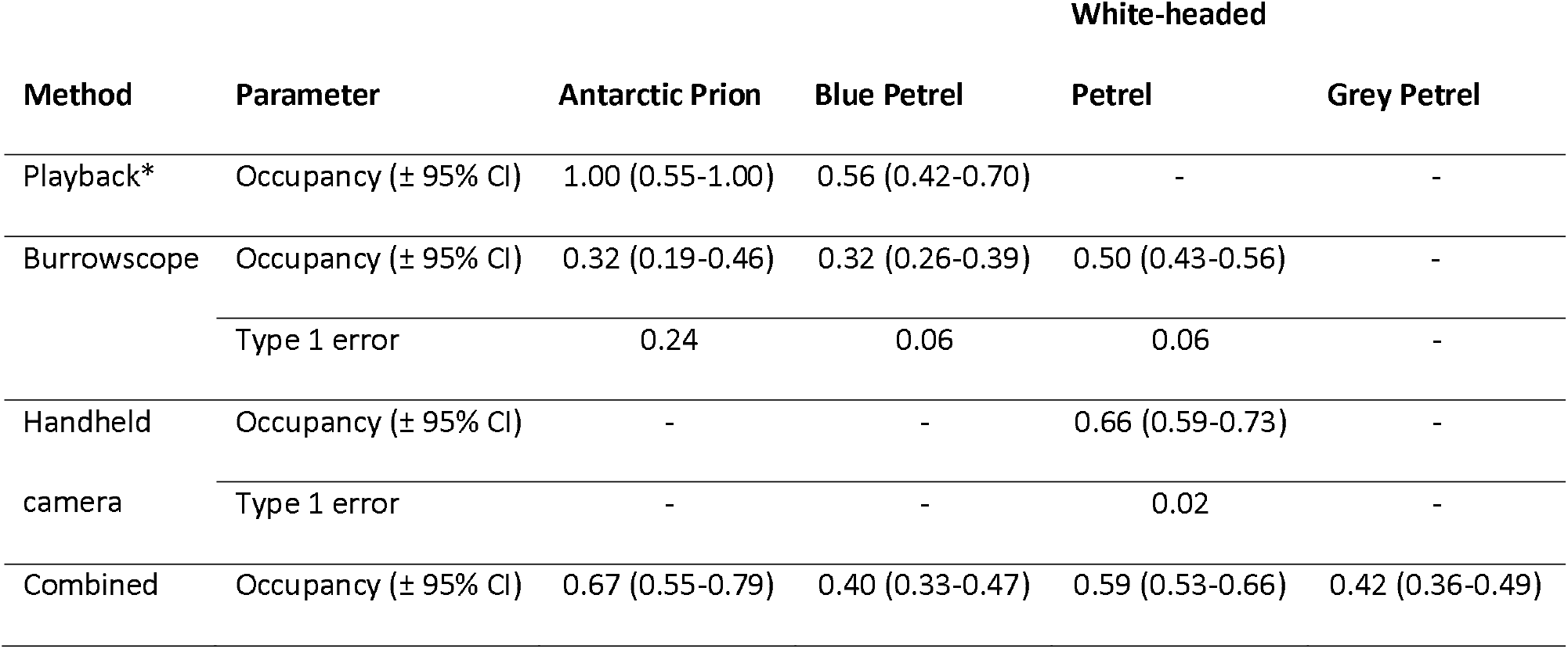
Burrow occupancy in different species estimated using different survey methods. Occupancy and confidence intervals are calculated from diagnosed burrows only. Type 1 error was not calculated for playback because occupancy status of all non-responsive burrows was listed as ‘uncertain’ owing to the possibility of non-responsive birds.

Uncertainty was lowest when using the burrowscope at larger White-headed Petrel burrows. We were able to correctly diagnose 218 of 233 (92%) burrows on the first visit, but 15 occupied burrows (6%) were incorrectly diagnosed as empty. Uncertainty was greater with a handheld camera than a burrowscope (25% compared with 2%, Figure 6) but type 1 error was not (Table 4).

Comparing methods for determining breeding status and breeding success, it is clear that playback is incompatible with collecting the burrow-level information needed. Even with manual burrow inspection, we were often unsure of breeding status on first visits alone because birds were not manoeuvred to check whether they were incubating eggs in order to avoid undue disturbance. However, by repeating visits through the season, we were able to determine the breeding status of 96 of 136 (70%) occupied White-headed Petrel burrows and 82 of 101 (82%) occupied Grey Petrel burrows (Table 5). We estimated breeding success as 0.83 (0.75-0.90) and 0.77 (0.68-0.75) for White-headed and Grey Petrels respectively after all visits. The two visits that were made to Blue Petrel burrows were insufficient to estimate breeding success.

**Table 5:**
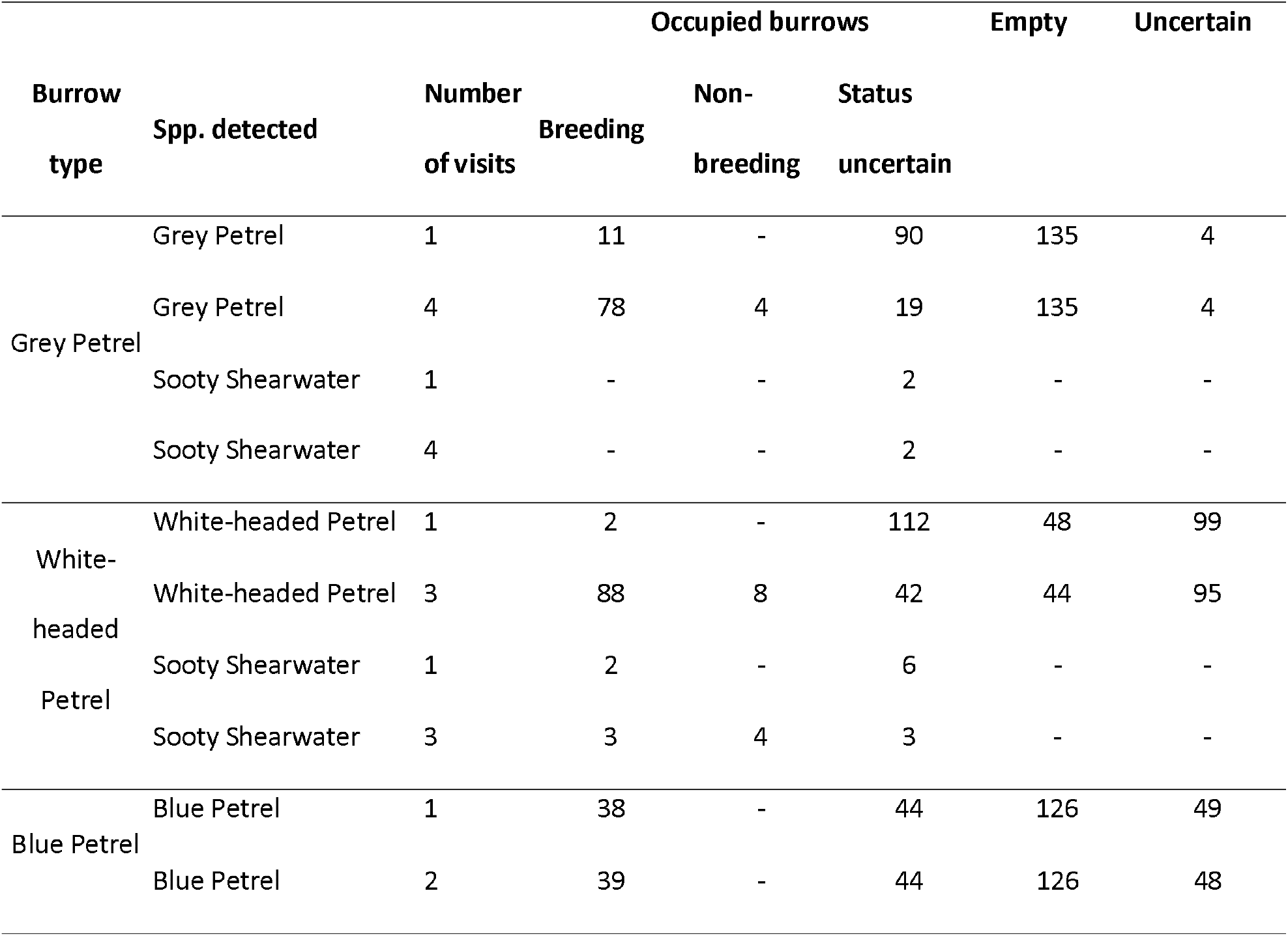
Inferred breeding status of seabird burrows based upon initial, and follow-up visits.

| Burrow<br>type | Spp. detected | Number<br>of visits | Breeding | Occupied burrows |  | Empty | Uncertain |
| --- | --- | --- | --- | --- | --- | --- | --- |
|  |  |  |  | Non-breeding | Status<br>uncertain |  |  |
| Grey Petrel | Grey Petrel | 1 | 11 | - | 90 | 135 | 4 |
|  | Grey Petrel | 4 | 78 | 4 | 19 | 135 | 4 |
|  | Sooty Shearwater | 1 | - | - | 2 | - | - |
|  | Sooty Shearwater | 4 | - | - | 2 | - | - |
| White-headed<br>Petrel | White-headed Petrel | 1 | 2 | - | 112 | 48 | 99 |
|  | White-headed Petrel | 3 | 88 | 8 | 42 | 44 | 95 |
|  | Sooty Shearwater | 1 | 2 | - | 6 | - | - |
|  | Sooty Shearwater | 3 | 3 | 4 | 3 | - | - |
| Blue Petrel | Blue Petrel | 1 | 38 | - | 44 | 126 | 49 |
|  | Blue Petrel | 2 | 39 | - | 44 | 126 | 48 |

Occupancy is an inferior metric to breeding status of burrows but is used in one-off survey visits. Through deployments (long and short) of camera traps at burrow entrances we believe it is possible to reliably determine burrow breeding status given a sufficient sample size of known-status burrows from which to model status as a function of nest-level activity. Furthermore, camera traps can substantially reduce uncertainty in estimated breeding success.

## Discussion

Conservation assessments focus on the portion of populations that are breeding, i.e. those which carry the reproductive potential of the species (IUCN, 2019). The challenge of determining the size and trends of burrowing-seabird breeding populations when only burrow entrances are visible at the surface is now decades old, yet there has been little research into the activity of seabirds at burrow entrances or the relationship between this activity and breeding status and success of individual burrows. We found that camera traps positioned at burrow entrances were able to collect time series activity patterns from which we could infer breeding status and breeding success of individual burrows. Provided the cameras used are fit-for-purpose, camera traps provided higher resolution data on the number of successful breeding attempts from a sample of burrows than is possible through burrow inspection.

Despite our relatively low sample size, we were able to discriminate effectively between breeding and non-breeding burrows using partial time-series data, suggesting that a thorough investigation of type 1 and 2 error rates around inferred breeding status is warranted. LDAs had higher sensitivity when identifying breeding burrows than non-breeding burrows from our small test dataset. Activity patterns associated with breeding are likely to be relatively uniform given that incubation shift duration and provisioning rates are all energetically determined (Chaurand and Weimerskirch, 1994; Jouventin, 1985). In contrast, non-breeding birds are not bound to a chick and can take longer absences. Non-breeding shearwaters are known to spend up to four times longer at the surface than breeding birds (Bonnaud et al., 2009), consistent with our observation of the highest activity peaks outside non-breeding burrows and the group means from LDAs.

Importantly the ratio of breeding to non-breeding burrows varies in dynamic populations. Populations undergoing expansion, for example following invasive species eradication, can have high proportions of immigrant and immature birds which may prospect without attempting to breed. In declining populations experiencing high predation rates prospecting birds can be disproportionately impacted resulting in a higher proportion of occupied burrows supporting breeding birds (Bonnaud et al., 2009). Differentiating between breeding and non-breeding burrows is, therefore, important, but it is difficult with traditional methods which we show are prone to high uncertainty. For future studies we would focus on increasing our sample size to assess whether linear discriminant models can be improved to increase their predictive ability. It would also be valuable to see how variable burrow entrance activities are between breeding sites. Calibration might only be necessary at a handful of sites to train LDAs that can be applied elsewhere to new camera trap datasets.

Regardless of the eventual reliability of camera trap LDAs for inferring breeding status of burrows, using camera traps in consort with other methods provides valuable information. For example, in the non-breeding burrows we followed, the camera traps revealed high rates of burrow activity but paired with manual inspection we found these burrows were rarely occupied in the daytime. Playback studies suffer from the untested assumption that occupied burrows support breeding birds (Schulz et al., 2006). Camera trap deployments could potentially be used to test this assumption and calibrate playback for use as a rapid assessment method. Additionally, the activity data from camera traps could be used to inform survey design by providing information on peak seasonal activity. Interestingly the early chick rearing LDA from mid-season was best at predicting breeding status of our test burrows. Most surveys focus on the start of the breeding season when birds are most vocal and responsive to playback and burrow inspection causes low disturbance or the end of the breeding season when breeding success can be inferred.

Camera traps have a number of potential advantages over traditional survey methods. Playback is only applicable to occupancy estimation, not to studying breeding numbers and success. We have found burrow inspection to be observer biased, its reliability varies according to the quality of the burrow-scope available (Lavers et al., 2019), and it can cause disturbance. The accuracy of estimates of breeding success are also dependent upon the frequency of burrow checks, whereas camera traps can yield precise timings of key events such as fledging or predation. Camera trap studies are less susceptible to personnel bias, and once established can potentially be repeated with relative ease and minimal disturbance. It is unclear how representative an occupancy estimate is when it is based upon data from a single season as is often the case. Occupancy varies between years because some species do not breed annually (Chastel, 2008), and stochastic events make some years more conducive to breeding than others. Adopting methods that can span several breeding seasons with little inter-annual bias in data collection will help to uncover some of these important patterns in seabird breeding dynamics.

We recorded several unexpected species and behaviours at cameras with serendipitous benefits. There is no reliable data on historic seabird populations on Macquarie Island before the impacts of invasive species (Brothers, 1984). Small petrels were reportedly abundant on coastal slopes but their identity is uncertain (Jones, 1980). Within living memory Antarctic Prions have been confined to the upland plateau so our camera trap images of them frequenting a lowland Blue Petrel colony and entering Blue Petrel burrows are the first records of the species utilising lowland areas since pest eradication. Similarly, White-headed Petrels have a restricted range and were reportedly absent from the north of the island, however, one individual at a Grey Petrel burrow on North Head, the extreme northern end of Macquarie Island and outside their current breeding range was recorded on our cameras. While these observations highlight the potential of cameras for gathering ecological data, Fischer et al.’s (2017) conclusion that camera traps are unsuitable for studying breeding biology at individual diving-petrel nests also applies in part to our study species. Not all entries and exits are captured by camera traps, and the precise position of the deployed camera can strongly affect what is recorded. Camera traps would, therefore, not be reliable for high resolution behavioural and phenological research such as studying incubation shift duration, and the frequency of chick provisioning.

We did encounter a number of challenges that will need addressing if camera traps are to be adopted more widely in burrowing seabird studies. Most important was the vast gulf in performance between the two makes of camera we used. Technologies are improving all the time, with open source platforms now allowing remote sensing to be taken to scale for major monitoring programmes (Hill et al., 2019), but it is worth noting the importance of budgeting appropriately for survey equipment when the costs associated with visiting seabird islands are so high. Second, tagging images is difficult. Citizen science platforms and improvements in machine learning are transforming the ability of monitoring programmes to tag large image datasets (Norouzzadeh et al., 2018; Willi et al., 2019), but behaviours may still be problematic. We had difficulty differentiating chicks from adults close to fledging. Only by manually coding images did we develop a familiarity with individuals which could aid identification. For example, bills of procellariform petrels develop slowly, and chicks often had noticeably slender bills even close to fledging, but judgements like this are highly subjective and potentially beyond a classification algorithm or volunteer programme. In one case it became apparent that two different chicks from adjacent burrows were triggering one camera. They were at different stages of development and could be readily identified. Ultimately this allowed us to confirm fledging success and timing from two burrows the status of which was uncertain based upon manual burrow-inspection, but after the first bird fledged the second switched burrows temporarily. If this behaviour is common it adds an additional challenge to determining breeding success. Cameras also needed some maintenance during the season. As well as battery and SD card replacement, some lenses developed condensation, and vegetation growth affected false trigger rates and obscured images.

We have shown that camera traps are highly effective at recording nest-level activity and behaviours of burrowing seabirds, and importantly that breeding status can be modelled as a function of these activity patterns. Significant investment is needed in terms of financial and human resources to purchase cameras, deploy and maintain them, calibrate camera data with burrow breeding status, and to tag images. Many of these investments can be front-loaded and we argue are worthwhile for long-term monitoring thanks to the benefits camera traps deliver—reduced uncertainty, widely applicable methodologies that can be applied by non-technical field workers, and low impacts on study species. For short surveys this effort is unlikely to be worthwhile and traditional surveys will be necessary.

## Authors’ contributions

JB conceived the ideas and designed methodology; JB and PP collected the data; JB analysed the data; JB led the writing of the manuscript. All authors contributed critically to data interpretation, drafts and gave final approval for publication.

## Acknowledgements

The authors thank Noel Carmichael and Tasmania Parks and Wildlife Service for their support facilitating this project. Thanks to Calum X. Cunningham and Toby D. Travers for suggests for data manipulation and analysis. This study was supported by funding from the Australian Government’s National Environmental Science Program through the Threatened Species Recovery Hub, the Australian Antarctic Science program (AAS 4305. JB was supported by a Research Training Program scholarship, an Antarctic Science International Bursary, National Environmental Science Programme Threatened Species Recovery Hub Research Support and a BirdLife Australia Stuart Leslie Bird Research Award. All methods were approved by the University of Queensland Native/Exotic Wildlife and Marine Animals (NEWMA) animal ethics committees (AE29713), the Macquarie Island Research Advisory Group and the Department of Primary Industries, Parks, Water and Environment (TFA 17305). Access to Macquarie Island was granted by the Tasmania Parks and Wildlife Service (Access Authority No. 17-18 5).

## Data Sharing

Raw data and full code for the analysis are available from the Australian Antarctic Data Centre.

