## Supporting Information for "Using camera traps to determine occupancy and breeding in burrowing seabirds"

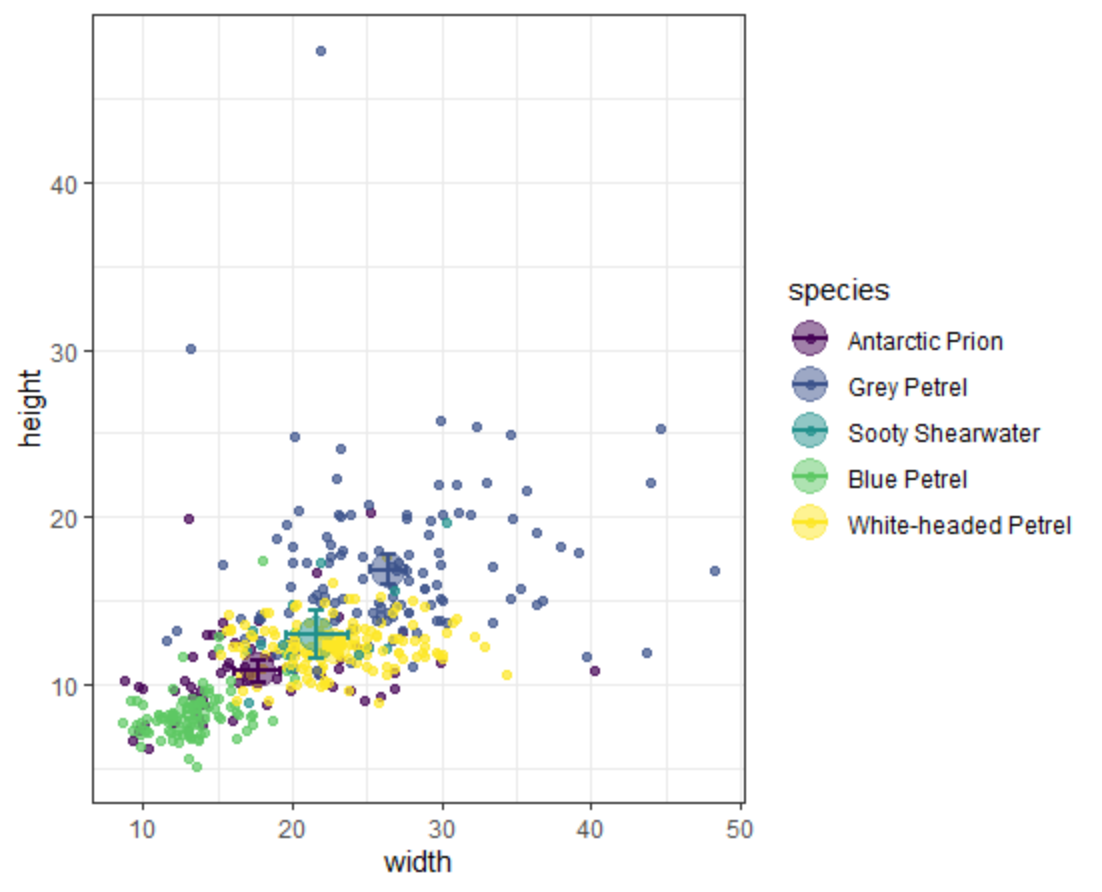

**Figure S1:** Dimensions of occupied seabird burrows on Macquarie Island. Larger circles represent the mean ± 95% CI. Despite overlap in dimensions, burrows were readily identified by their general impression, size and shape.
